# Cell recovery and DNA extraction methods skew relative abundance estimations of foodborne pathogens and spoilage organisms on stainless steel surfaces

**DOI:** 10.1101/2025.11.03.686371

**Authors:** Jingzhang Feng, Sarah E. Daly, Katerina Roth, Abigail B. Snyder

**Author notes:** **Names and e-mail addresses for all authors**: Jingzhang Feng, Sarah E. Daly, Katerina Roth, Abigail B. Snyder. **Contact information for Corresponding Author**: Abigail B. Snyder.

## Abstract

Studies investigating surface microbiota in food facilities often include estimates of relative abundance. However, obtaining accurate relative abundance values can be challenging. Differences among microbes can lead to different degrees of cell recovery from the surface and different DNA extraction yields that skew downstream relative abundance estimates. Here, we evaluated (1) the impact of cell recovery from surfaces using sponge swabs and (2) DNA extraction protocol on the relative abundance estimates of relative abundance on artificially inoculated surfaces. Our results showed that *Escherichia coli* (Gram-negative cell)*, Listeria monocytogenes* (Gram-positive cell), *Bacillus cereus* (bacterial spore), *Alicyclobacillus suci* (bacterial spore), *Exophiala phaeomuriformis* (fungal cell), *Aspergillus fischeri* (fungal spore) differed significantly (p<0.05) in their recovery rates from stainless steel surfaces, ranging from 2.9%±3.0% recovery to 94.9%±3.0% recovery. Modification of the DNA extraction protocol by extending the bead-beating step by 10 min generally improved DNA yields, though the impact varied by organism. For example, DNA yields of *E. coli* increased from 70 to 84 ng/mL while that of *L. monocytogenes* increased only from 23.2 to 29.2 ng/mL. Amplicon sequencing results indicated that the differences in cell recovery and DNA extraction among microbial species skewed the relative abundance estimates from inoculated surfaces. For example, the estimated relative abundance of *L. monocytogenes* was 9-17%, which was lower than its actual relative abundance (25%). These results underscore the limitations of surface microbiota characterization in food facilities and highlighted the need to improve current recovery and DNA extraction methods.

**Importance:** Amplicon sequencing has been used to characterize microbial communities on facility surfaces. However, few studies have evaluated the accuracy of the amplicon sequencing workflow for quantifying spoilage and pathogenic organisms in these microbial communities. Here, we assessed the accuracy for amplicon sequencing to evaluate the relative abundance of spoilage and pathogenic organisms commonly found in food processing environments. The results revealed biases in relative abundances due to limitations in cell recover and DNA extraction methods. These findings revealed the potential biases in surface microbiota characterization in food facilities, and the need to refine current recovery and extraction methods to enhance the accuracy of microbiota characterization.

## 1. Introduction

Amplicon sequencing has been used to characterize microbial communities on food facility surfaces (1–5). This process involves recovering cells from environmental surfaces with swabs, extracting DNA from the recovered cells, amplifying targeted DNA regions with Polymerase-Chain-Reaction (PCR), and sequencing the DNA amplicons (6). Nucleotide sequences are then mapped to databases (*e.g.* NCBI RefSeq) to determine the taxonomy of organisms within the microbiota (7). Subsequently, the relative abundance of each taxon is calculated to reflect its proportion within the community (8). The relative abundances of spoilage (e.g. *Alicyclobacillus suci, Exophiala phaeomuriformis, Aspergillus fischerii*) and pathogenic (e.g. *Escherichia coli, Listeria monocytogenes, Bacillus cereus*) organisms are often compared among samples from various environmental conditions (9). Food producers are especially interested in characterizing and tracking the harborage points of pathogenic bacteria and spoilage organisms to assess and prevent product contamination risks.

Food facility surfaces contain diverse spoilage and pathogenic organisms including Gram-positive and Gram-negative bacteria, yeasts, and molds. Due to differences in physiochemical properties (*e.g.* sizes, cell wall thickness), these organisms exhibit different surface attachment, which affect their removal from surfaces during recovery (10–12). As a result, different proportions of the cells of different microbes may be recovered from the surface via swabbing, causing the estimations of their relative abundances to be skewed (13–15). While some studies have compared the recovery of Gram-positive and Gram-negative bacteria using sponge swabs, differences in recovery among other cell morphologies have not been examined (13, 16). Therefore, it remains unclear to what extent sponge swabs can introduce recovery biases that skew relative abundance calculations.

Further bias in relative abundance calculations can be introduced during DNA extraction. The distinct physicochemical properties of different cell types can impact DNA extraction and result in inconsistent DNA yields. Specifically, the structural differences between organisms can impact cell integrity, which affects the efficiency of lysis and DNA release. This issue is particularly prominent in spore-forming organisms, since they contain a hard cell barrier that resist lysis (17, 18). However, commonly used DNA extraction methods (*i.e.* DNA PowerSoil Kit) can produce low DNA yields for spore-forming organisms (17, 19). Consequently, the relative abundance of spore-formers could be underestimated when using this method.

Additionally, any modified methods need to maintain consistent DNA yield across organisms to prevent biases. Although some optimized methods have been proposed, little research has assessed the impact of these modified methods on relative abundances. Furthermore, few investigations have been carried out to evaluate method performance on spoilage organisms that can form spores in food facilities.

In summary, issues with recovery and extraction yields can lead to inaccurate assessment of relative abundances of relevant spoilage and pathogenic microbes on surfaces in the food industry. This study aims to evaluate the effects of the sponge swab recovery method and modified DNA extraction protocols on the relative abundances of spoilage and pathogenic organisms (**Figure 1**). Understanding the impact of these methods can inform strategies to improve accuracy and understand the limitations of microbiota characterization.

**Figure 1.**
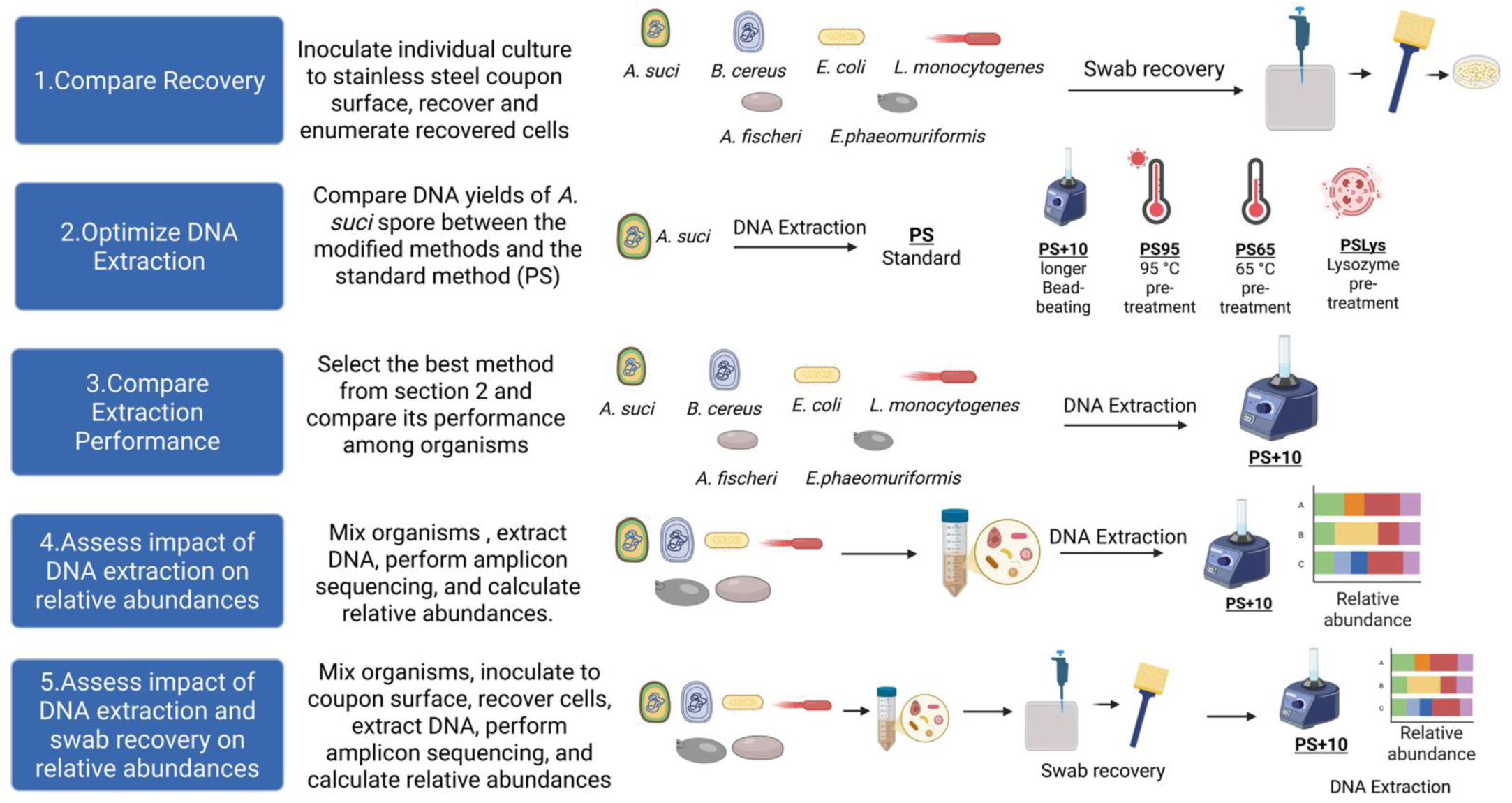
Workflow to evaluate the impact of recovery and DNA extraction on downstream amplicon sequencing to assess microbes from surfaces in food processing environments.

## 2. Materials and methods

### 2.1 Creating spore and vegetative cell cultures

#### 2.1.1 Creating Alicyclobacillus suci and Bacillus cereus spore stocks

*Alicyclobacillus suci* FSL W10-0048 was grown on Yeast-Starch-Glucose (YSG) (pH 4.0) agar at 45°C for 7days to allow sporulation (20). *Bacillus cereus* FSL A1-001 was grown on Brain-Heart-Infusion (BHI) agar at 35°C for 24hrs, then an isolated colony was transferred to AK#2 agar (BD, Thermo Fisher Scientific, Waltham, MA, USA) and incubated at 35°C for 7days to allow sporulation. *A. suci* and *B. cereus* spores were harvested by adding 1 mL of sterile DI water and gently scraping the surface of the plate with a plate spreader. The remaining vegetative cells were removed by treatments of 200mg/mL lysozyme at 37°C for 40minutes, 1% Sodium Lauryl Sulfate at 30°C for 20minutes, then an 80°C heat shock for 10minutes. The presence of spores in the final suspension was confirmed by malachite green staining. Spore concentrations were confirmed by cell counts on hemacytometer and colony counts of heat shocked cultures (80°C for 10minutes) on agars (YSG for *A. scui*, BHI for *B. cereus*).

#### 2.1.2 Exophiala phaeomuriformis cells and Aspergillus fisherii ascospores collection

*E. phaeomuriformis* and (FSL E2-0572) *A. fisherii* (NRL B-2354) were grown separately on Acidic-Potatoes-Dextrose-Agar (APDA) (pH 4.0) and incubated at 25°C for 28days. Colonies were harvested with 1xPBS and pelleted. Pellets were washed and resuspended again in 1xPBS (21). The presence of fungal cells and ascospores in the final suspension was confirmed by phase contrast microscopy (AMScope, Irvine, CA, USA). Fungal culture concentrations were confirmed by the colony counts on APDA and cell counts using a hemacytometer.

#### 2.1.3 Escherichia coli & Listeria monocytogenes cultures

Both *E. coli* FSL A1-0127 and *L. monocytogenes* FSL B2-0079 were incubated separately in Trypticase Soy Broth at 35°C for 24hrs. Vegetative cells were pelleted, washed, and resuspended in 1x PBS. Concentrations of vegetative cultures were confirmed by colony counts on BHI agar and cell counts using a hemacytometer.

### 2.2 Assessing recovery rate of each cell culture from stainless steel coupon surfaces

Each cell culture was adjusted to approximately 8log CFU/mL, serially diluted and plated on agars to confirm its concentration. Ten drops (10µL per drop) of the adjusted culture were inoculated onto a 1ft x 1ft sterilized stainless-steel coupon (7log CFU/coupon in total). Stainless steel coupons were dried under a fume hood for 20minutes. Cells on stainless-steel coupon surfaces were collected by 3M swabs (Neogen, Lansing, MI, USA) premoistened with 10mL of Dey-Engly broth (Lahou and Uyttendaele, 2014). The handle of the 3M swabs was snapped off and 10 mL of 1x PBS was added to the sponge. The sponge was stomached at 260rpm for 2minutes. Cell pallets were collected by transferring sponge buffers to a 50mL falcon tube and centrifuge at 6500xg for 15minutes. Cell pellets were resuspended in 1mL of 1× PBS. One hundred microliters of each resuspended culture were serially diluted and plated on BHI to enumerate *B. cereus*, *E. coli*, and *L. monocytogenes*, plated on YSG to enumerate *A. suci*; and plated on APDA to enumerate *E. phaeomuriformis* and *A. fischerii*. BHI plates were incubated at 35°C for 24hrs, YSG plates were incubated at 45°C for 48hrs, and APDA plate were incubated at 25°C for 72hrs. The recovery percentage for each culture was determined using the following equation:

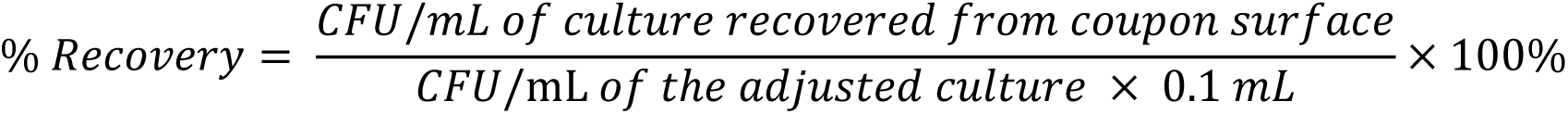

### 2.3 Determine the optimized method for DNA extraction on A. suci

Four modified DNeasy PowerSoil Pro kit (Qiagen, Redwood City, CA, USA) protocols and the standard PowerSoil kit protocol (**PS**) were tested by extracting DNA from 5log CFU/mL of *A. suci* spores. The cell concentration of the spore culture was confirmed using a hemacytometer prior to extraction. The modifications include: 1) the standard protocol with an additional 10 minutes of beads beating (**PS+10**), 2) heat pre-treatment at 95°C for 10minutes in CD1 buffer (**PS95**), 3) heat pretreatment at 65°C for 10minutes in CD1 **PS65**), 4) enzymatic pretreatment in lysis buffer containing 10mg/mL lysozyme plus 5mg/mL proteinase K at 37°C for 10minutes (**PSlys**) (3). Each DNA extraction method was performed in triplicate. Extraction yields were quantified based on nucleic acid concentration (ng/mL) in the lysate using a Qubit fluorometer (Thermo fisher scientific, Waltham, MA, USA). Ct values of 16S V3–V4 amplicons from each lysate were determined by Quantitative-Polymerase-Chain-Reactions (qPCRs). Specifically, 2µL of each lysate was added to a master mix containing 12.5µL of SYBR Green PCR Master mix (Thermo fisher scientific, Waltham, MA, USA), 1µL each of 10µM forward and reverse primer, 8.375µL of ddH_2_O, and 0.125µL of ROX control. qPCR was conducted in QuantiStudio 6 (Thermo fisher scientific, Waltham, MA, USA) with the following conditions: Initial denaturation at 95°C for 2minutes, 40 cycles of: 95°C for 5seconds, 60°C for 10seconds.

### 2.4 Evaluation of modified DNA Extraction methods on other spores and vegetative cells

The PS+10 method demonstrated improved DNA yield from *A. suci* spore extractions and was therefore selected to evaluate extraction efficiency across all cultures. Suspensions with 5log, 4log, and 3log CFU/mL of each bacterial culture, and suspensions with 6log, 5log, 4log, and 3log CFU/mL of each fungal culture were prepared. The cell concentration of each culture was confirmed by the cell count under hemacytometer. All suspensions were pelleted prior to extractions. DNA was extracted from pellets using either the PS or PS+10 method. DNA yields were quantified by measuring DNA concentrations from 5log CFU/mL of bacterial cultures and 6log CFU/mL of fungal cultures using Qubit. Extracts from cultures with lower cell concentrations were below the quantification limit and were therefore not quantified.

### 2.5 Extract DNA from microbiota

Three types of microbiota were prepared, each containing two bacterial spore cultures (*i.e. A. suci*, *B. cereus*), two vegetative cell cultures (*E. coli* and *L. monocytogenes*), and two fungal cultures (*A. fischeri* and *E. phaeomuriformis*). One microbiota contained an equal ratio of each culture. The second type of microbiota contained 6log CFU/mL of each bacterial spore and fungal culture and 5log CFU/mL of each vegetative cell culture, representing a high spore-to-vegetative cell ratio. The third type of microbiota contained 5log CFU/mL of each bacterial spore and fungal culture, and 6log CFU/mL of each vegetative cell culture, representing a low spore-to-vegetative cell ratio. Cell concentrations were confirmed by hemacytometer counts prior to creating the microbiota. Triplicates of each microbiota were prepared. DNA from each microbiota was extracted with method PS+10. Extraction lysates were sent to NovoGene co. (Beijing, China) for 16S v3-v4 and ITS2 amplicon sequencing.

### 2.6 Assessing the amplification rate of 16s V3-V4 region for bacterial DNA and ITS 2 regions for fungal DNA

One milliliter of each bacterial and fungal culture was pelleted by centrifuging at 12,000xg for 5minutes. Each pellet was subjected to DNA extraction using the PowerSoil kit, and the resulting lysates were quantified with Qubit to determine DNA concentration. The DNA concentration of each lysate was adjusted to 200ng/mL with ddH_2_O. Two microliters of each adjusted lysate were added to the qPCR master mix and performed qPCR with conditions described in **section 2.3**.

### 2.7 Recovering microbiota from surfaces and performing amplicon sequencing

Triplicates of microbiota with 6log CFU/mL of cells per culture were created. Cell concentrations were confirmed by hemacytometer prior to creating the microbiota. Microbiota cultures were centrifuged at 12,000xg for 5minutes to pellet cells. Pellets were resuspended in 100µL of 1x PBS, inoculated onto coupons, recovered with swabs, and pelleted as described in **section 2.2**. DNA was extracted from the pallet using the PS+10 method. Extraction lysates were sent to NovoGene co. (Beijing, China) for 16S v3-v4 and ITS2 amplicon sequencing.

### 2.8 Assessing the impact of swabbing on relative abundance

Viable cells of each bacterial and fungal species in the microbiota described in **section 2.7** were enumerated before inoculation and after recovery using selective media to assess the impact of swab recovery on relative abundances. For enumeration, 100µL of serially diluted culture was plated onto YSG to enumerate *A. suci*, on VRBG to enumerate *E. coli*, on LMPN to enumerate *L. monocytogenes*), and on APDA to enumerate *E. phaeomuriformis* and *A. fischeri*. An additional 100 µL aliquot was heat-shocked at 80°C for 10min, serially diluted, and plated on BHI to enumerate *B. cereus*. Incubation conditions were: BHI and LMPN at 35°C for 24h; VRBG at 37°C for 24h; APDA at 25°C for 72h; and YSG at 45°C for 48h. Relative abundance of each viable bacterial or fungal organism before inoculation and after recovery was calculated as follows:

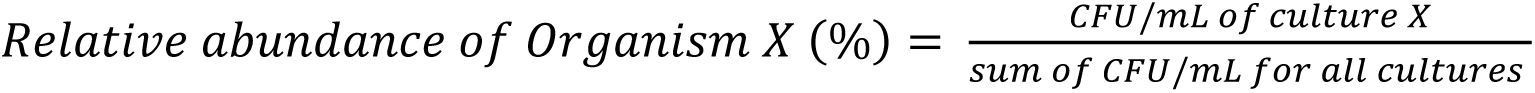

### 2.9 Sequence Analysis

Raw sequencing data was processed with QIIME 2 (22.2) (v. qiime-2022.2) (23). Sequences were quality checked with DADA2 (24). Paired end reads were denoised, had chimeras removed, and were trimmed if their quality score was below 15. A feature table of Amplicon-Sequence-Variants (ASVs) was generated after denoising and was analyzed by a pre-trained classifier based on the SILVA 16S v3-v4 database to create a taxonomy table (25). The abundance of each taxon was normalized by its 16S rRNA copy number using the q2-gcn-norm plug-in in QIIME 2. Microbial profiles were generated, and the relative abundance of each microbe was calculated with the phyloseq package in R (version 1.44.0) (26).

### 2.10 Statistical analysis

Significant differences in recovery among cell culture types were determined by ANOVA followed by Tukey’s post-hoc test. Differences in extraction yields and Ct values between the modified methods and the standard methods were evaluated by paired-end t-tests. Differences in relative abundances between sequencing results and theoretical values (***i.e.*** calculated based on cell counts using a hemacytometer) were assessed using t-tests with Bonferroni correction.

## 3. Results and discussion

### 3.1 A lower proportion of vegetative bacterial cells was recovered through surface swabbing than bacterial spores, fungal vegetative cells, and fungal ascospores

The percent of recovered *L. monocytogenes* and *E. coli* were similar. Although the recovery of *L. monocytogenes* was numerically higher than *E. coli*, the difference was not statistically significant (p>0.05) (**Table 1**). On average, 2.9%±3.0% of the inoculated *E. coli* cells, and 6.6%±6.0% of the inoculated *L. monocytogenes* cells were recovered from surfaces. Both percentages were lower than those reported by Keeratipibul et al., 2017 (*E. coli*: 52.3±6.0%, *L. monocytogenes*: 73.7%±2.6%) but the *L. monocytogenes* recovery was similar to that observed by Gómez et al., 2012 (*L. monocytogenes*: 2.81±0.63%) (28, 29). The reason for the high recovery percentages reported by Keeratipibul et al. (2017) may be due to the use of log-transformed numbers which inflates recovery percentages.

**Table 1.** The amount (log CFU/ft^2^) and proportion of the inoculated viable cells recovered from stainless steel surfaces with sponge swabbing.

| Kingdom | Species | Morphology | Gram Reaction | Average Inoculated Cells ± Standard Deviation (log CFU/ft<sup>2</sup>) | Average Recovered Cells ± Standard Deviation (log CFU/ft<sup>2</sup>) | Average % Recovery ± Standard Deviation<sup>1</sup> |
| --- | --- | --- | --- | --- | --- | --- |
| Bacteria | *E. coli* | Vegetative | Negative | 7.25±0.03 | 5.49±0.54 | 2.9%±3.0%<sup>a</sup> |
| | *L. monocytogenes* | Vegetative | Positive | 7.7±0.21 | 6.34±0.48 | 6.6%±6.0%<sup>a</sup> |
| | *B. cereus* | Spores | Positive | 6.82±0.12 | 6.8±0.12 | 94.9%±5.0%<sup>b</sup> |
| | *A. suci* | Spores | Positive | 6.21±0.34 | 5.96±0.46 | 35.3%±11%<sup>c</sup> |
| Fungi | *E. phaeomuriformis* | Yeast-like conidia | NA | 6.91±0.12 | 6.82±0.24 | 80%±19%<sup>b</sup> |
| | *A. fischerii* | Ascospores | NA | 5.51±0.09 | 4.72±0.61 | 36%±12%<sup>c</sup> |

<sup>1</sup>Values followed by same letters are not significantly different (p>0.05)

Notably, the number of cells that died-off on surfaces during drying could also bias the recovery percentages. Specifically, cells susceptible to desiccation, such as *E. coli* and *L. monocytogenes,* may have a lower recovery percentage due to their die-off. However, studies by Wilks et al., 2005 and Wilks et al., 2006 reported a reduction of less than 1 log CFU of *E. coli* and *L. monocytogenes* cells on stainless steel surfaces within 180 min of incubation at room temperature (30, 32). Since cells were dried for less than 30 min in this study, the number of dead cells is unlikely to substantially affect the recovery percentage of *E. coli* and *L. monocytogenes*.

The recovery of other bacteria differed significantly from each other (**Table 1**). On average, sponge swabs recovered 94.9%±5.0% of *B. cereus* spores and 35.3%±11% of *A. suci* spores from surfaces. While significantly (p<0.05) higher than the two vegetative cells tested, these numbers were also significantly different from each other. Similarly, the recovery percentages of the two fungi were significantly (p<0.05) higher than those of vegetative cells and differed from each other. On average, 80%±19% of *E. phaeomuriformis* conidia and 36%±12% of *A. fischeri* ascospores were recovered from surfaces.

Differences in recovery between cell morphologies could be due to variations in physiological properties. These variations can affect the attachment of cells to surfaces, thus impacting their removal by swabs (33). Previous studies have identified cell size as a physiological property affecting cell removal (34, 35). For instance, Cai et al. (2021) demonstrated that fluid shear stress had a greater impact on cell removal when cell size exceeded the width of surface roughness topology (36). Specifically, the study showed that shear stress from rotational fluid had a greater impact on the removal of *E. phaeomuriformis*, a spore exceeding the roughness measurements of the surface, than on the removal of *L. monocytogenes* from stainless steel surfaces (36). This finding was consistent with our observation that sponge swabs recovered a lower percentage of *L. monocytogenes* cells from stainless steel surfaces than *E. phaeomurifomis* (**Table 1**). In fact, the effect of cell size may have also contributed to the recovery difference between the tested vegetative cells (*e.g. L. monocytogenes*) and bacterial spores (*e.g. A. suci*), since the tested bacterial spores (1.5-1.8 by 0.9-1.0 µm) had greater cell size than tested vegetative cells (0.5 by 2 µm) (37–39).

In addition to cell size, other physio-chemical factors, such as chemical composition and hydrophobicity, had also been identified as factors that could affect cell removal (10–12). Importantly, these physiological properties can also affect cell adhesion to swabs, leading to different release capacities for different organisms (10, 40). Differences in release can further contribute to the swab recovery difference (13). Consequently, swab recovery difference can affect analyses linking organism abundances to environmental characteristics. Understanding factors impacting cell recovery using swabs can inform strategies to improve recovery. For instance, recognizing that hydrophobicity of cell membranes can affect removal, some methods have added surfactants to swabs to improve cell recovery (40). Enhanced recovery could provide a more accurate understanding of the association between microbiota characteristics and environmental attributes.

### 3.2 Extending the duration of bead beating improved DNA yields for spores

Because of the variability in starting concentration among trials, the results in **Figure 2** were shown individually for each trial. Extending beating time (PS+10) increased DNA yield consistently across all three trials (Trial 1: +2.4 ng/mL, Trial 2: +0.6 ng/mL, and Trial 3: +4.8 ng/mL) compared to the standard method (**Figure 2A**). A significant (p<0.05) increase in DNA yield using the PS+10 method was confirmed by a paired-end t-test. This result aligned with the previous finding that extending bead-beating time by 10 min increased extraction yield from *Campylobacter jejuni* by 1.6-1.8 folds (41). On the other hand, the method with 95°C heat pre-treatment (PS95), 65°C heat pre-treatment (PS65) and lysozyme pretreatment (PSlys) did not consistently improve extraction yields, although some trials produced higher DNA yields than the standard method. Quantitative Polymerase Chain Reaction (qPCR) was conducted to evaluate the amplification rate of each extraction product in downstream PCR (**Figure 2B**). None of the modified methods (PS+10, PS65, PS95, and PSLys) consistently lowered Ct values, suggesting that these modifications did not enhance amplification efficiency. Interestingly, while the PS+10 method increased DNA yields, it did not improve amplification rates. The Ct value lowered slightly in two trials (Trial 1: -0.39, Trial 3: -0.46) but increased in one trial (Trial 2: +1.01) when using PS+10. This outcome could be attributed to DNA shearing caused by extended bead beating. Excessive mechanical force produced from a longer bead beating could fragment DNA molecules released form spores (42, 43). The fragmented DNA may not be amplified in PCR as readily and therefore the Ct value did not improve (44). PCR is a key step that enables downstream sequencing and analysis to determine identity, taxonomic classifications, and relative abundance calculations (45). Hence, while the PS+10 method increased DNA yield, it may provide limited improvements to the accuracy of relative abundance assessments for spores. Future studies should investigate strategies to reduce DNA shearing due to bead beating (**e.g.** increase bead beating intensity but shorten the duration). These strategies could further improve the accuracy of relative abundance assessments in microbial community analysis (46)

**Figure 2.**
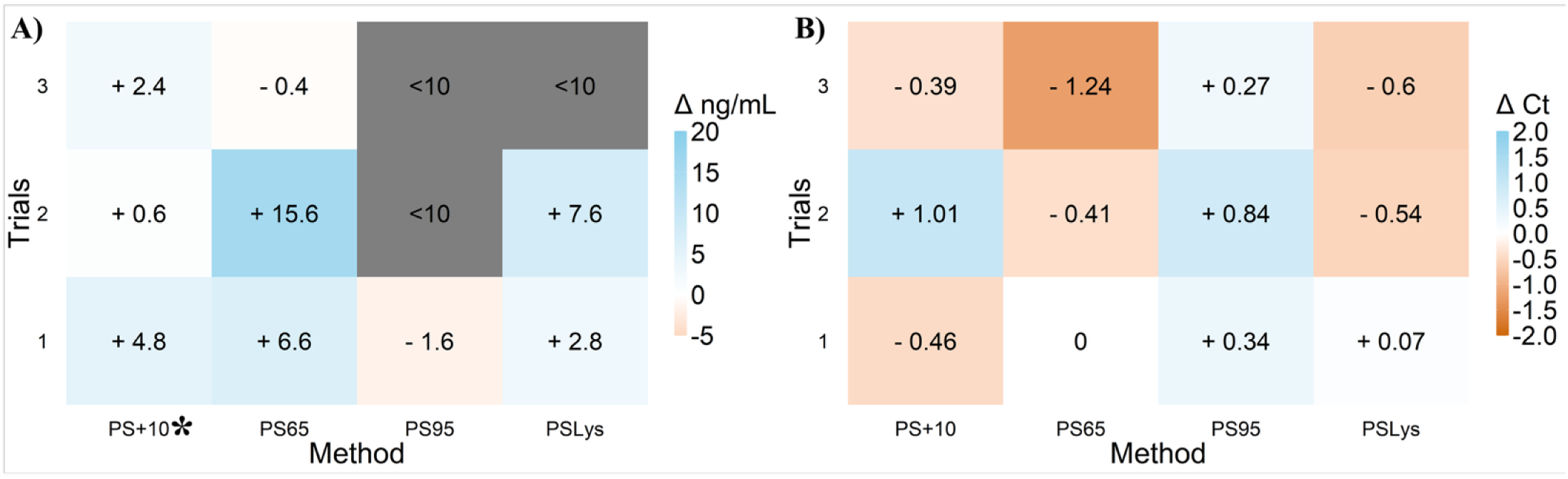
Changes to the DNA yield (**A**) and Ct values generated by qPCR (**B**) compared to the standard method (PS) when using the modified extraction methods (PS+10, PS65, PS95 or PSLys). Asterisks (*) indicates the modified method with significant increase in DNA yields compared to the standard method. Gery color grids indicate that results were below the detection limit of Qubit (<10 ng/mL).

### 3.3 The PS+10 method did not improve extraction yield uniformly across different bacteria and fungi

The PS+10 method improved DNA yields for some but not all organisms (**Figure 3A & B**). The extended beads beating method (PS+10) significantly (P<0.05) increased DNA yields from *A. suci, B. cereus, E. coli,* and *E. phaeomriformis*. PS+10 increased the extraction yield of *E. coli* by 14 to 40 ng/mL, which is the greatest among all the tested organisms. Besides *E. coli,* PS+10 also increased the extraction yield of *A. suci* by 6.2 to 8.4 ng/mL, and *B. cereus* by 5.4 to 6.4 ng/mL. Among fungal organisms, PS+10 significantly (P<0.05) improved DNA yield only from *E. phaeomuriformis*, with increases ranging from 125 to 567 ng/mL. In contrast, PS+10 did not improve DNA extraction from *L. monocytogenes* and *A. fischeri*. For *L.* monocytogenes, PS+10 slightly increased DNA yield (3.0ng/mL) in trial 3 but reduced yields in trial 1 (-5.0 ng/mL) and trial 2 (-3.2 ng/mL). Similarly, PS+10 increased the extraction yield of *A. fischeri* in trial 1 (+2 ng/mL) and 3 (+ 254 ng/mL), but reduced yields in trial 2 (4 ng/mL). *L. monocytogenes* and *A. fischeri* have previously been identified as organisms with low DNA extraction efficiencies (47, 48). The lack of improvement in DNA yield for these organisms suggests that their relative abundance may be underestimated when applying the PS+10 method.

**Figure 3.**
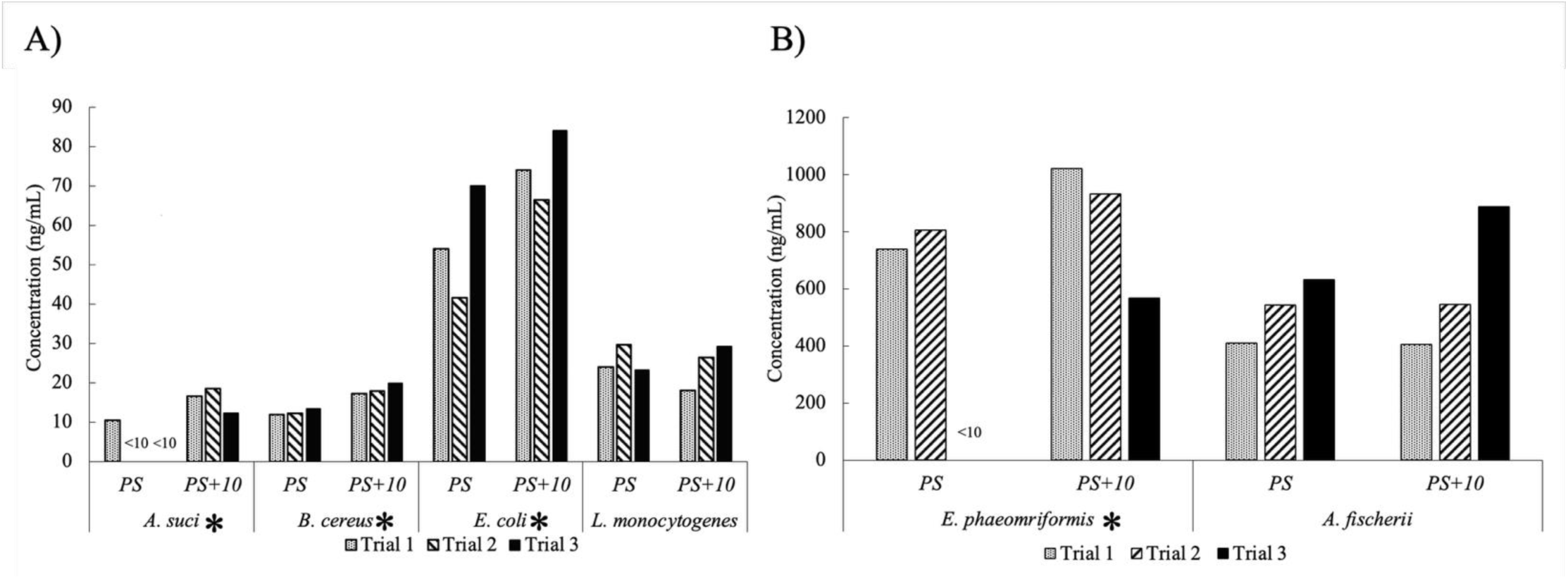
DNA extraction using the PS and PS+10 methods across the selected cultures. Extraction yields were compared between the standard method (PS) and the PS+10 method for each bacterial (**A**) and fungal (**B**) organism. Asterisk (*) indicates a significant (p<0.05) increase in extraction yield when using the modified method (PS+10).

The efficacy of the PS+10 method was likely affected by cell rigidity. This was evident by the high extraction efficiency of *E. coli* cell, which contains a permeable cell wall that facilitate lysis and DNA release (48, 49). In addition to rigidity, cell size could also affect extractions. Smaller cells are more difficult to recover by centrifugation during DNA extraction, resulting in lower DNA yields (50). This may explain why PS+10 exhibited lower extraction efficiency for *L. monocytogenes* (0.5 by 2 µm) than for *A. suci* (1.5-1.8 by 0.9-1.0 µm) and *B. cereus* (1.6 by 0.9 µm) spores, despite all having rigid cell structures (37–39). Differences in the extraction efficiency of PS+10 could skew the relative abundance of bacterial and fungal organisms. Therefore, better protocols are needed to enhance extraction efficiency for cells with small sizes or rigid structures, such as *L. monocytogenes*.

### 3.4 The PS+10 method skewed the relative abundance of organisms

The observed relative abundances of organisms were significantly (P<0.05) different from the theoretical values (**Figure 4A**). The only exception was the relative abundance of *A. suci* in the microbiota with a high spore low vegetative cell ratio, in which 38.9%, 39.3%, and 43.8% of *Alicyclobacillus* was observed in trial 1, 2 and 3 respectively. These percentages were similar (p>0.05) to the theoretical value (45.5%). Differences between the theoretical and observed relative abundances indicated that using PS+10 in DNA extraction did not produce a result that accurately reflected the proportion of organisms in the microbiota. Furthermore, we consistently recorded higher observed relative abundances of *Escherichia* and *Exophiala* than their theoretical relative abundances across all three ratios tested. For instance, in trial 1, the relative abundance of *Escherichia* was 70.6%, 37.3%, and 70.2% in microbiota with equal, high, and low spore-to-vegetative cell ratios, and the relative abundance of *Exophiala* was 82.1%, 94.2%, and 86.7% in the respective ratios. These percentages were all significantly (P<0.05) greater than their respective theoretical relative abundances (*Escherichia:* 25.0%, 4.55%, and 45.0%, *Exophila:* 50% for all three microbiota). The overestimation of *Escherichia* and *Exophiala* relative abundances aligned with the elevated extraction efficiency of these genera as shown in **Figure 3**. This suggested that the discrepancy in extraction efficiency may have contributed to the bias of relative abundance. In fact, a similar trend has been observed in organisms with lower relative abundances. For instance, the underestimation of *Bacillus* across microbiota corresponds to its lower extraction efficiency shown in **Figure 3**.

**Figure 4.**
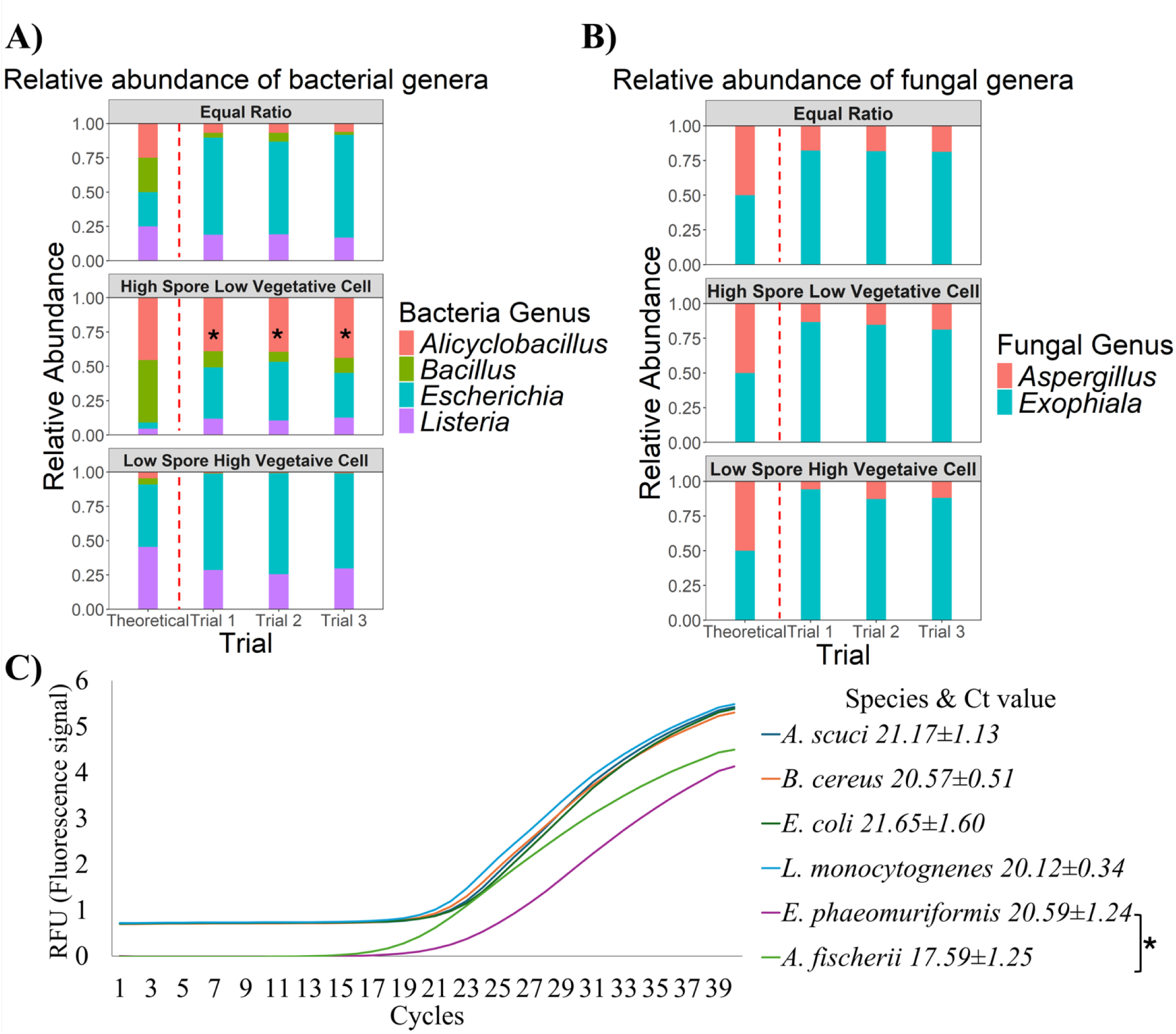
The genus level relative abundances of bacterial (**A**) and fungal (**B**) organisms as a result of extracting DNA from three ratios of microbiota using the PS+10 method. These ratios include equal ratio of bacterial spores and vegetative cells, high ratio of bacterial spore low ratio of vegetative cell, and low ratio of bacterial spore high ratio of vegetative cell. The theoretical relative abundances were calculated according to the cell counts through hemacytometer of each organism in the microbiota. Fungal genera had equal theoretical relative abundances across all microbiota since equal concentrations of each fungal organism were included in the mixtures.

The qPCR results further suggested that DNA extraction, rather than amplification via PCR, was the primary contributor to the skewed relative abundances. The Ct values generated by qPCR were used to assess the relative amplification efficiency targeting the 16S V3-V4 region or the ITS2 region for each organism. A lower Ct value indicates a higher amplification rate, which can lead to an over-estimated relative abundance for the corresponding organism (51). However, the Ct values of bacterial organisms were at a similar (p>0.05) level ranging from 20.12±0.34 to 21.65±1.60, indicating that differences in amplification rates did not contribute substantially to the biases of relative abundance (**Figure 4C**). Although amplification rates differed significantly between *E. phaeomuriformis* (20.59±1.24) and *A. fischeri* (17.59±1.25), these differences did not align with the observed differences in their relative abundances. Specifically, *A. fischeri* had a higher amplification rate than *E. phaeomuriformis,* yet the observed relative abundance of *A*.

*fischeri* was consistently lower than its theoretical value. This suggested that discrepancies in PCR amplification rates may not be the major cause of relative abundance biases for fungal organisms. Together, these results implied that biases in relative abundances were mostly attributable to DNA extraction.

Incomplete extractions of cells with rigid structures have been identified as the major factor in skewing relative abundance calculations (52). Previous studies have suggested that increasing the extraction efficiency of hard-to-lyse cells is key to enhancing the accuracy of relative abundances (49, 53). Although the PS+10 method increased DNA yields from bacterial spores, the improvement was not sufficient to enhance the accuracy of relative abundance assessments. These findings further highlight the need to optimize the extraction efficiency for hard-to-lyse cells. In addition, the reduced quality of DNA due to the extended bead-beating treatment may be another factor negatively affecting relative abundances. As a result, developing methods to minimize DNA shearing during extractions may improve the accuracy of relative abundance assessments.

Asterisk (*) indicates the genus with similar (p>0.05) relative abundance as the theoretical value. qPCR was conducted to assess the PCR amplification efficiency for each organism, targeting the 16S V3-V4 or ITS 2 region using equal concentrations of DNA from each organism (**C**). Asterisk (*) indicates a significant (p<0.05) difference in amplification rates between organisms.

### 3.5 Recovery with sponge swabs followed by DNA extraction with the PS+10 method caused changes to relative abundances and microbiota composition

Sample collection, processing, DNA extraction and analysis affected the relative abundance of bacterial genera (**Figure 5A**). After following these processes, the sequencing derived relative abundances of *Listeria* (trial 1: 12%, trial 2: 9%, trial 3: 17%) were significantly (p<0.05) lower than the theoretical value (25%), whereas those of *Bacillus* (trial 1: 37%, trial 2: 44%, trial 3: 36%) were significantly (p<0.05) higher. On the other hand, the relative abundances of *Alicyclobacillus* (trial 1: 22%, trial 2: 24%, trial 3: 21%) and *Escherichia* (trial 1: 29%, trial 2: 23%, trial 3: 26%) were similar to theoretical values. Interestingly, the process of sample collection, processing, DNA extraction and analysis resulted in a different bias to relative abundance variation between the sequencing derived and theoretical values, compared to extraction alone (**Section 3.4**, **Figure 4**). For example, in **Figure 4**, the relative abundances of *Bacillus* in the sequencing result were lower than the theoretical value, whereas in **Figure 5A**, it was higher. Differences among sequencing results suggest that both swab recovery and DNA extraction contributed to the skewed relative abundances compared to theoretical values.

**Figure 5.**
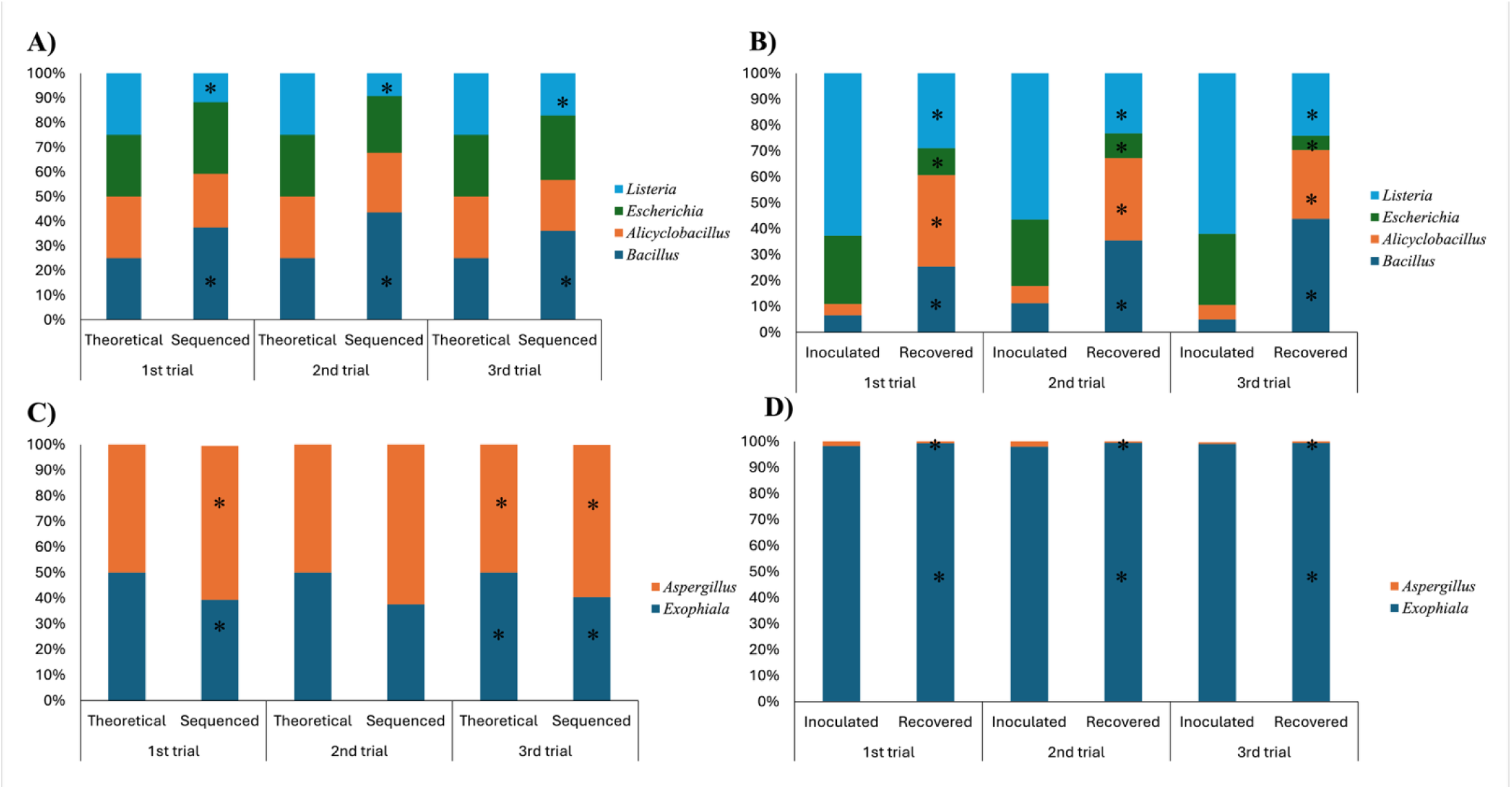
Relative abundances (**A**) changes of bacteria organisms following cell recovery, DNA extraction, and amplicon sequencing of microbiota inoculated onto stainless steel surfaces at equal cell ratios. The theoretical relative abundances were calculated according to the cell counts using a hemacytometer. Asterisk (*) indicates the organism with significantly (P<0.05) different relative abundance compared to the theoretical value. Surface microbiota was also plated on selective media and enumerated. The proportions of each bacterium in the inoculated and recovered microbiota were determined from colony counts and compared (**B**). Asterisk (*) indicates the organism with significant (P<0.05) different relative abundance between the inoculated and recovered microbiota. The same analyses and comparisons were made between the fungal communities (**D & E**).

To specifically assess the impact of swab recovery, we quantified viable cells of each bacterial species using selective media and compared counts between the inoculated and recovered microbiota. The results indicated that bacterial relative abundances differed between the inoculated and recovered microbiota (**Figure 5B**). These changes also corresponded with the recovery efficiency of individual organisms (**Table 1**). For instance, *E. coli*, an organism with low recovery efficiency (2.9% ± 3.0%), showed a decrease from the inoculated level of ∼26% to 10% in trial 1 and trial 2, and 5% in trial 3 based on the recovered microbiota. Conversely, *B. cereus*, an organism with high recovery efficiency (94.9% ± 5.0%), showed an increase in from the inoculated level of 7% to 25% in trial 1, 11% to 35% in trial 2, and 5% to 44% in trial 3.

These trends confirmed that recovery partially contributed to the skewed relative abundance observed in sequencing results. Notably, *L. monocytogenes* was underestimated due to the impact of DNA extraction (**Figure 3&4**) and swab recovery (**Figure 5A & Table 1**). Consequently, food safety and quality concerns associated with these organisms may be challenging to address when evaluating surface microbiota with amplicon sequencing (54, 55). Therefore, research aimed at improving swab recovery and DNA extraction should place emphasis on pathogens such as *Listeria*.

The relative abundance of the fungal community also changed following sample recovery, DNA extraction, and analysis (**Figure 5 C**). The sequencing derived relative abundances of fungal organisms were significantly (P<0.05) different from those based on theoretical composition (**Figure 5C**). The relative abundances of *E. phaeomuriformis* (trial 1: 39%, trial 2: 38%, trial 3: 40%) in the sequencing results were significantly lower (P<0.05) than the theoretical value (50%), while those of *A. fischeri* (trial 1: 60%, trial 2: 63%, trial 3: 60%) were significantly higher (P<0.05) (**Figure 5C**). When comparing viable cell concentrations in the inoculated and recovered microbiota, we observed a significantly higher (P<0.05) relative abundance of *E. phaeomurifomis* and a significantly lower (p<0.05) lower relative abundance of *z. fischeri* via plating (**Figure 5D**). The relative abundance of *E. phaeomuriformis* increased from 98.0% to 99.9% in trial 1, 98.0 to 99.5% in trial 2, and 99.0% to 99.4% in trial 3 between the inoculated and recovered microbiota. In contrast, *A. fischeri* decreased from 2.00% to 1.00% in trial 1, 2.00% to 0.50% in trial 2, and 1.00% to 0.60% in trial 3 between the inoculated and recovered microbiota. It remains unclear why the calculated relative abundance of *E. phaeomuriformis* from amplicon sequencing was lower than the theoretical value, even though biases introduced by both recovery (**Figure 5D**) and DNA extraction (**Figure 4B**) were expected to overestimate its abundance. One possible explanation is that current bioinformatic analysis pipelines (*e.g.* DADA2), which are primarily designed for 16S rDNA gene analysis, can generate errors when clustering fungal ITS sequences into taxonomic units (56, 57). These errors may have caused the sequencing derived values to be different from the theoretical value (58–60). Potential errors in ITS sequence analysis highlight the need to improve bioinformatic methods for analyzing fungal ITS data.

## 4. Conclusion

Amplicon sequencing analysis distorted the relative abundance of spoilage and pathogenic organisms. This bias was attributed to differences in cell recovery and DNA extraction among microbes. Variations in physiological properties and cell integrity may contribute to these differences. To improve the accuracy of relative abundance estimates, future studies should investigate cellular mechanisms affecting recovery and DNA extraction and develop strategies to mitigate their impact, particularly for the pathogen *Listeria monocytogenes*. Mechanisms associated with Gram-positive organisms need greater attention due to their lower recovery and DNA extraction efficiency. Additionally, method optimizations are needed to enhance the analysis of fungal sequencing data. Such efforts will improve relative abundance estimates and enable more accurate identification of key microbiota using amplicon sequencing.

### CRediT authorship contribution statement

**Jingzhang Feng:** Writing-original draft, Visualization, Methodology, Investigation, Data curation, Conceptualization. **Sarah E. Daly:** Writing – review and editing. **Katerina Roth:** Writing – review and editing. **Abigail B. Snyder:** Writing – review and editing, Supervision, Project administration, Funding acquisition, Conceptualization.

### Data Availability

Data will be made available on request

### Declaration of Competing Interest

All of the authors have seen and approved the submitted manuscript, and there are no possible conflicts of interest.

## Acknowledgement

This research was supported by the U.S Department of Agriculture’s (USDA), National Institute of Food and Agriculture project 2022-67017-36289. The findings and conclusions in this preliminary publication have not been formally disseminated by the U.S Department of Agriculture and should not be constructed to represent any agency determination or policy.

